# Sign-epistatic centromere drive in panmictic populations

**DOI:** 10.1101/2022.10.24.513597

**Authors:** Evgeny Brud

## Abstract

Comparative work has revealed a highly non-random elevation of karyotypic homogeneity *within* mammalian species for either telocentric chromosomes or centric fusions, and a karyotypic bimodality *among* species for these binary chromosome morphologies. A verbal theory developed by Pardo-Manuel de Villena and Sapienza argues that morphology-biased segregation in female meiosis explains the corresponding directional evolution in favor of one or the other chromosome form within species, and moreover, periodic reversals of meiotic spindle asymmetry explain the pattern of bimodality observed among species. Here I investigate a population genetic model in which I assume that the direction of the spindle asymmetry is under the control of a modifier gene, either linked or unlinked to a focal karyotypic mutant (linkage being to a centric fusion, say), and I derive the corresponding invasion conditions for the modifier-centric-fusion gene complex. I demonstrate that the scenario put forth in the verbal theory can be explained by a two-step process in which (1) a centric-fusion invades to fixation while the linked modifier winds up at an intermediate frequency by hitchhiking, and then (2) subsequent fixations of unlinked centric fusions occur. Via numerical iteration of the model, I demonstrate that the typical post-hitchhiking frequency of the linked modifier (from step 1) is broadly sufficient for subsequent unlinked invasions (step 2). Sign-reversing modifier evolution is therefore concluded to be a plausible mechanism instantiating the principles of a female drive theory of karyotype evolution.

## 1. Introduction

Asymmetric female meiosis presents an opportunity for selfish centromeric variants to occupy a greater than 50% share of the eggs in heterozygous females.^1–3^ Recent work in *Mimulus* and *Mus* demonstrates that chromosomes can vary within species for centromere strength, a character defined by the abundance of centromeric histone H3 (CENP-A), the epigenetic marker of centromeres that recruits kinetochore proteins during mitotic and meiotic divisions.^4–6^ Females of *Mus musculus* with a heterozygous make-up of strong and weak centromeres exhibit non-Mendelian segregation in the first meiotic division, with stronger variants having a 60% probability of segregating to the secondary oocyte. A significant advance in understanding the mechanism of centromere drive was discovered by Akera et al. (2017), who demonstrated that meiotic spindles in *Mus musculus* are asymmetrically organized on the two poles with respect to levels of tyrosinated and detyrosinated alpha-tubulin, and so the two halves of the spindle have different stability properties.^7^ Later work showed that destabilizing molecules are recruited preferentially to the cortical pole (leading to the first polar body) by the expanded kinetochore^25^ (see [26] for recent review). Heterozygosity for centromere strength therefore results in an elevated rate of spindle detachment of the unequal bivalent during metaphase. Stronger centromeres in *Mus musculus* showed a marked elevation in their rates of detachment from the cortical pole as compared to the egg pole, followed by preferential re-orientation toward the egg.

Earlier comparative work predicted spindle asymmetries in female meiosis as an explanation of macroevolutionary patterns of karyotype evolution.^8^ Female drive was argued to underlie the observation that the majority of mammals have karyotypes that are either mostly acrocentric (functionally one-armed; here not distinguished from telocentric) or mostly metacentric (bi-armed), with relatively few species carrying an even mixture of both chromosome morphologies. As argued by Pardo Manuel de Villena and Sapienza^3^, the simultaneous realization of three asymmetry principles in female trivalent segregation (unequal centromere numbers, an asymmetrical meiotic spindle, and the alternative fates of meiotic products) causes the spread of either acrocentrics or metacentrics within a species karyotype. In this case, differential centromere strength would owe to differences in centromere number across a trivalent (1 vs. 2). Results consistent with a signature of centromere drive were observed in non-mammalian taxa as well, including ray-finned fishes^9^ and holocentric^10^ insects, as well as in studies of B chromosome patterns.^11–13^ Experimental work in the karyoptic races of Central European *Mus musculus* also implicates population-specific correlations between centromere strength and chromosome form, so that centromere size differences, in addition to centromere number differences, is a causative factor in the nonrandom segregation of female Robertsonian translocation heterozygotes.^5^

Reiterating: (i) predominance of a specific chromosome form would be expected to result from a meiotic spindle that favored a particular centromeric morphology (e.g. acrocentric) in female meiosis I, and (ii) the observation of frequent switches from one morphology to another would result from the reversal of the relevant spindle asymmetry during the course of evolution. ^8,24,26^ The verbal theory and the mammalian comparative pattern was presented in two articles by Pardo-Manuel de Villena and Sapienza (2001a,b). However, a corresponding mathematical model has not previously been worked out.

Here I investigate one genetically-explicit interpretation of the theory: I consider what happens in a two-locus model in the simultaneous presence of a centromere driver and a dominant modifier that governs the direction of the segregation ratio as mediated by an asymmetric spindle. Because the sign of the segregation advantage of a centromere variant depends on the make-up of the modifier locus, the model investigates the effect of *sign-epistatic centromere drive*. Furthermore, since the spindle is an organism-level trait, the spindle modifier is assumed to have a sign-changing effect on segregation distortion at all chromosomes in the genome. I ask: (1) What conditions favor the simultaneous rise in frequency of a karyotype variant and a linked spindle modifier that both start at low frequency? and (2) Does the aftermath of the ‘modifier-linked’ phase of karyotype evolution select for subsequent changes in the rest of the genome (i.e. at unlinked chromosomes)?

## 2. Model

Consider a two-sex random mating population of infinite size that varies at loci A and M for alleles A_1_, A_2_ and M_1_, M_2_, respectively. Locus A is a centromere and allelic variation at A refers to the presence of the unfused (A_1_) and fused (A_2_) state (i.e. A1 are telocentrics and A_2_ is a metacentric for a particular set of homologous chromosome arms). A_1_A_2_ heterozygotes exhibit unequal segregation ratios *k* A_1_: *1-k* A_2_ in M_1_M_1_ females, but the reverse segregation ratio in M_1_M_2_ and M_2_M_2_ females; segregation in males is Mendelian throughout. Male heterozygotes at A experience a constant reduction in fitness due to viability or fertility defects (independent of mating type) in all M genotypes. Assuming discrete non-overlapping generations, the system of recursions governing evolution is given by:

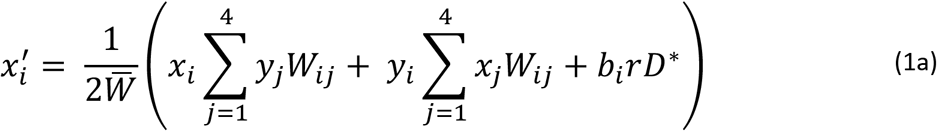

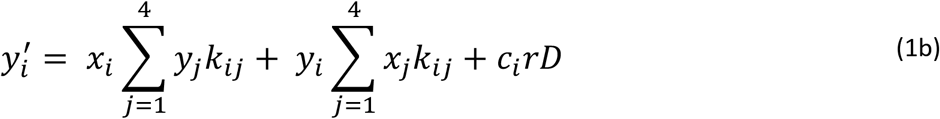

The equations, valid for i, j = 1, 2, 3, 4, track the evolution of gametic frequencies (A_1_M_1_, A_1_M_2_, A_2_M_1_, A_2_M_2_) in males (x_1_, x_2_, x_3_, x_4_) and females (y_1_, y_2_, y_3_, y_4_) in successive generations. The mean fitness in males, 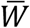, is given by the sum of all post-selection male genotype frequencies prior to normalization: 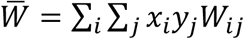. The gametic disequilibrium terms are *D* = *x*_1_*y*_4_ + *x*_4_*y*_1_ – *x*_2_*y*_3_ – *x*_3_*y*_2_ and *D*\* = (1 – *s*)*D*. The elements *W_ij_* and *k_ij_* refer to the elements in the matrices governing male fitness (**W**) and female segregation (**K**), respectively:

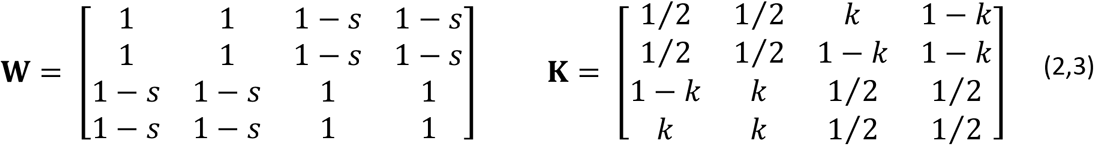

*b_i_* equals –1 when i = 1,4 and +1 when i = 2,3. *c*_1_ = – *k*), *c*_2_ = 1 – *k*, *c*_3_ = *k*, and *c*_4_ = (–1)*K*. I assume ½ ≤ *k* ≤ 1, 0 ≤ *s* ≤ 1, and 0 ≤ *r* ≤ ½ as bounds on parameter values.

## 3. Results

### 3.1 Step-one: modifier-linked drive of a centric fusion

#### 3.1.1 Double-invasion conditions

First I analyze the invasion dynamics at A1M1 haplotype fixation. The stability of the fixation equilibrium 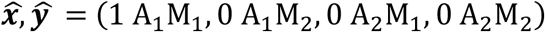 can be ascertained by analyzing the Jacobian matrix. Evaluating at 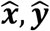, the Jacobian yields a 4 x 4 block matrix with elements that are 2 x 2 block matrices:

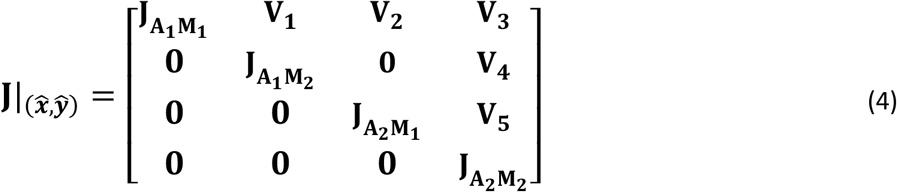

The stability of 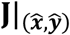 is determined by the spectra of the block matrices on the diagonal. **J_A_1_M_1__**, **J_A_1_M_2__** and **J_A_2_M_1__** correspond to resident population dyanmics in the absence of simultaneous rare variation for A_2_ and M_2_; these are never associated with an eigenvalue greater than one in magnitude. **J_A_2_M_2__** gives the invasion dynamics for simultaneous presence of A_2_ and M_2_, and yields the leading eigenvalue, 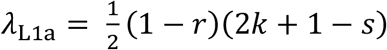.

##### Result 1a

λ_L1a_ is greater than one for the following space of parameter values (Fig. 1):

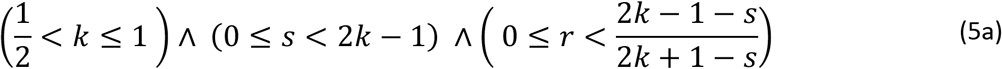

Or equivalently:

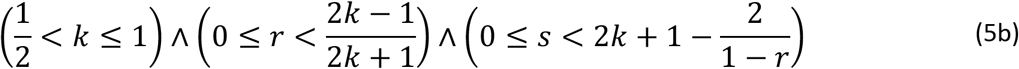

A mutant karyotypic driver (A_2_) and a linked sign-changing dominant modifier (M_2_) will both invade a population initially fixed for resident alleles, A_1_ and M_1_, as long as conditions 5a-b are satisfied with respect to the parameters of drive intensity (*k*), selection (*s*), and linkage (*r*).

**Fig 1.**
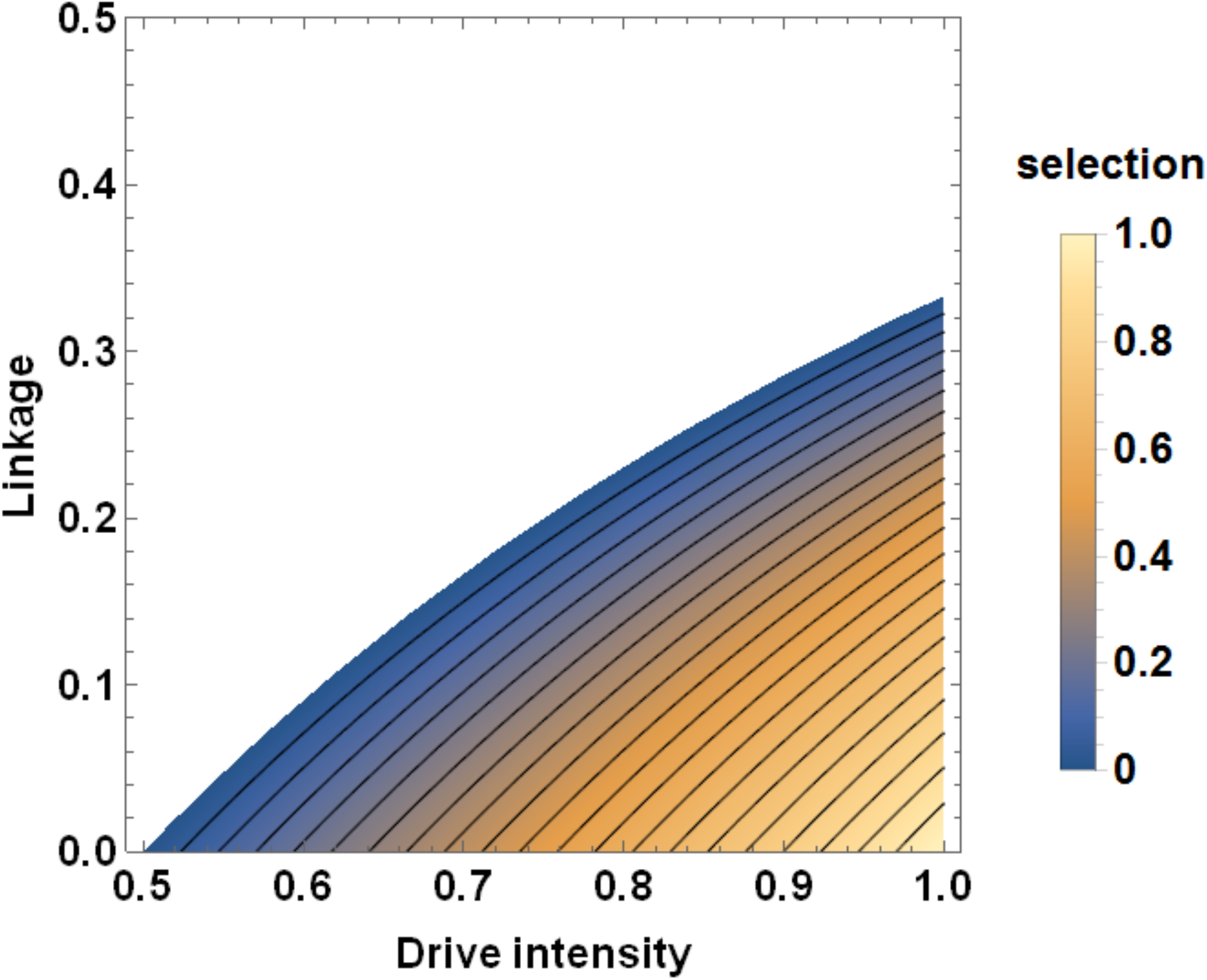
Invasible parameter region for the modifier-centric-fusion gene complex. The space of parameters from Result 1 are displayed (assuming pM_2,0_ = 0), with drive intensity in female meiosis on the horizontal axis, linkage on the vertical, and the selection coefficient of male underdominance given by the contours. The top right corner of the invasible region corresponds to linkage equal to 1/3.

The conditions on the recombination rate exclude the case of unlinked modifiers. Indeed no double-invasions involve *r* ≥ 1/3. As an example, for the case of 60% drive intensity with no male fitness reduction, the minimal recombination rate permitting double-invasion is ~0.09.

An extension of **Result 1a** to greater than infinitesimal frequencies of M_2_ is possible by adding a parameter pM_2,0_, which is the M_2_ initial frequency. Although the model under investigation is deterministic, one may imagine that in the absence of karyotypic variation, M_2_ is able to neutrally evolve toward higher rare frequencies than strictly zero (say, between 0% and 10%), at which point an A_2_ mutant is introduced at infinitesimal frequency. The generalization results in a Jacobian of the form^14^ 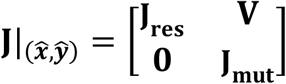, where **J_res_** is a 4 x4 block matrix corresponding to the resident population (leading eigenvalue never greater than one); **J_mut_** is a 4 x 4 block matrix that yields the leading eigenvalue:

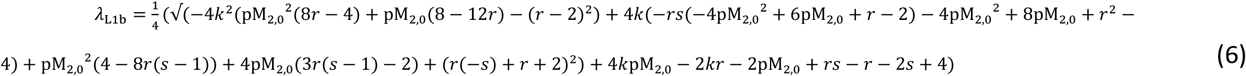

##### Result 1b

Assuming 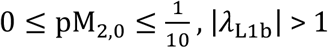 corresponds to the extended invasion conditions for A_2_M_2_:

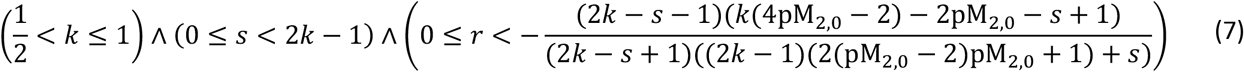

The right-hand side of the recombination rate inequality in expressions (5a) and (7) gives the maximal linkage (r_max_) consistent with mutant invasion; numerical cases show that small values of pM_2,0_ only modestly affect the value of r_max_, and so the extended conditions of **Result 1b** are well-approximated by **Result 1a**. For example, r_max_ attains its highest value when *k* = 1 and *s* = 0; for pM_2,0_ = 0 this value is 1/3 (as already reported above), while for pM_2,0_ = 1%, the corresponding r_max_ equals 0.34. For larger initial M_2_ frequency pM_2,0_ = 10%, r_max_ = 0.43, and so a substantial M_2_ presence is seen to moderately loosen the restrictions on permissible linkage. For k = 78% and s = 14%, the corresponding values for r_max_ are 0.1735, 0.1764, and 0.2094 for pM_2,0_ equal to 0%, 1%, and 10%, respectively. A broader look is achieved by comparing the total invasible regions of parameter space; this confirms overall insensitivity to small values of pM_2,0_ (Fig. S1).

Since no recessive deleterious effects are assumed in the model, an invading A_2_ driver increases until fixation, and linkage with the modifier causes a hitchhiking effect on M_2_. Because recombination acts to break down the association between alleles, M_2_ winds up at a neutral equilibrium frequency somewhere below fixation (Section 3.3).

#### 3.1.2 Conditions for monotonicity of invasion

The time-course of allele-frequency change in the model is sensitive to the initial level of gametic disequilibrium (D_0_). Even if the invasion conditions derived above (expressions 5, 7) are met, an insufficiently positive level of D_0_ results in negative frequency change of A_2_ for the first several tens or hundreds of generations (depending on parameter values), followed by a reversal in trajectory and the beginning of a rise to fixation. The cause of the trajectory-reversal owes to the build-up of positive D in the form of coupling gametes. A rare A_2_ driver which initially declines over many generations may reach biologically unrealistic allele frequencies (e.g. 10^-15^) prior to its increase. With sufficiently positive initial disequilibrium, the frequency changes for A_2_ and M_2_ are always positive. An expression for the minimal D (D_min_) consistent with a monotonic increase of mutant frequencies can be derived by reformulating the model in terms of D_0_ and the mutant allele frequencies at A (pA_2_) and M (pM_2_), and then setting the single-generation change in pA_2_ equal to 0, which yields a quadratic in D.

The biologically relevant root corresponds to

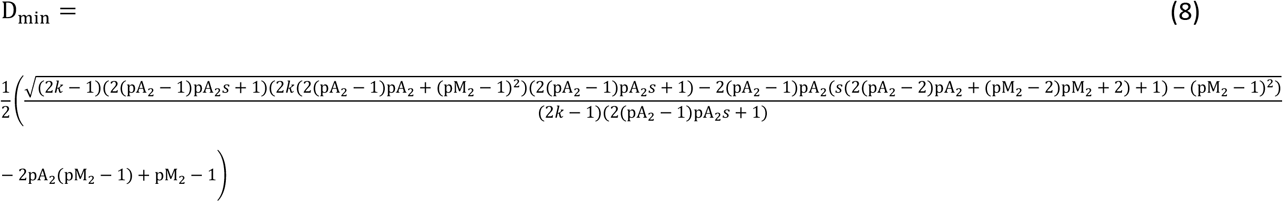

On the assumption that A_2_M_1_ is lower in frequency than A_1_M_2_ during the initial generation, a normalized version (i.e. Lewontin’s D’)^15^ is obtained by dividing D_min_ by the A_2_M_1_ frequency:

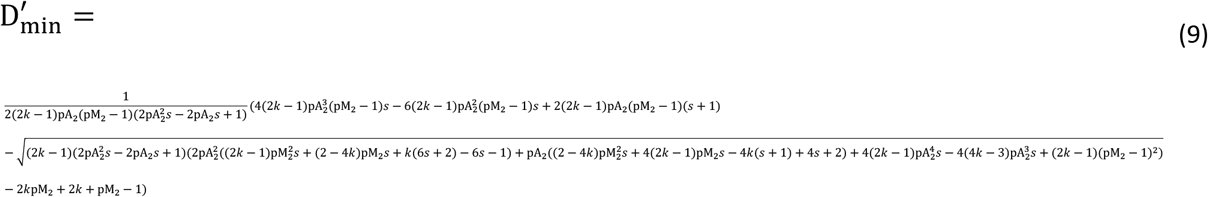

An initial value of D greater than D_min_ implies Δ(pA_2_) >0 in the first generations and thereafter. To avoid unrealistic trajectories as mentioned, restricting focus to D_0_ > D_min_ is sensible for numerical iteration (Section 3.3). Because 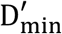, which always ranges from 0 to 1, is easier to interpret than the non-normalized D_min_, the monotonicity condition is most conveniently expressed as 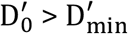.

For small values of the selection coefficient, 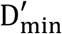 is relatively insensitive to changes in drive intensity, except near *k* = ½ (Fig. 2b,e). Neutral drive (*s* = 0, *k* > ½) with small pM_2,0_ gives 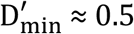 (Fig. 2a); an increase in the selection coefficient results in an increase to 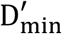 (Fig. 2c,f). (Notably, linkage does not figure in the expression).

**Fig 2.**
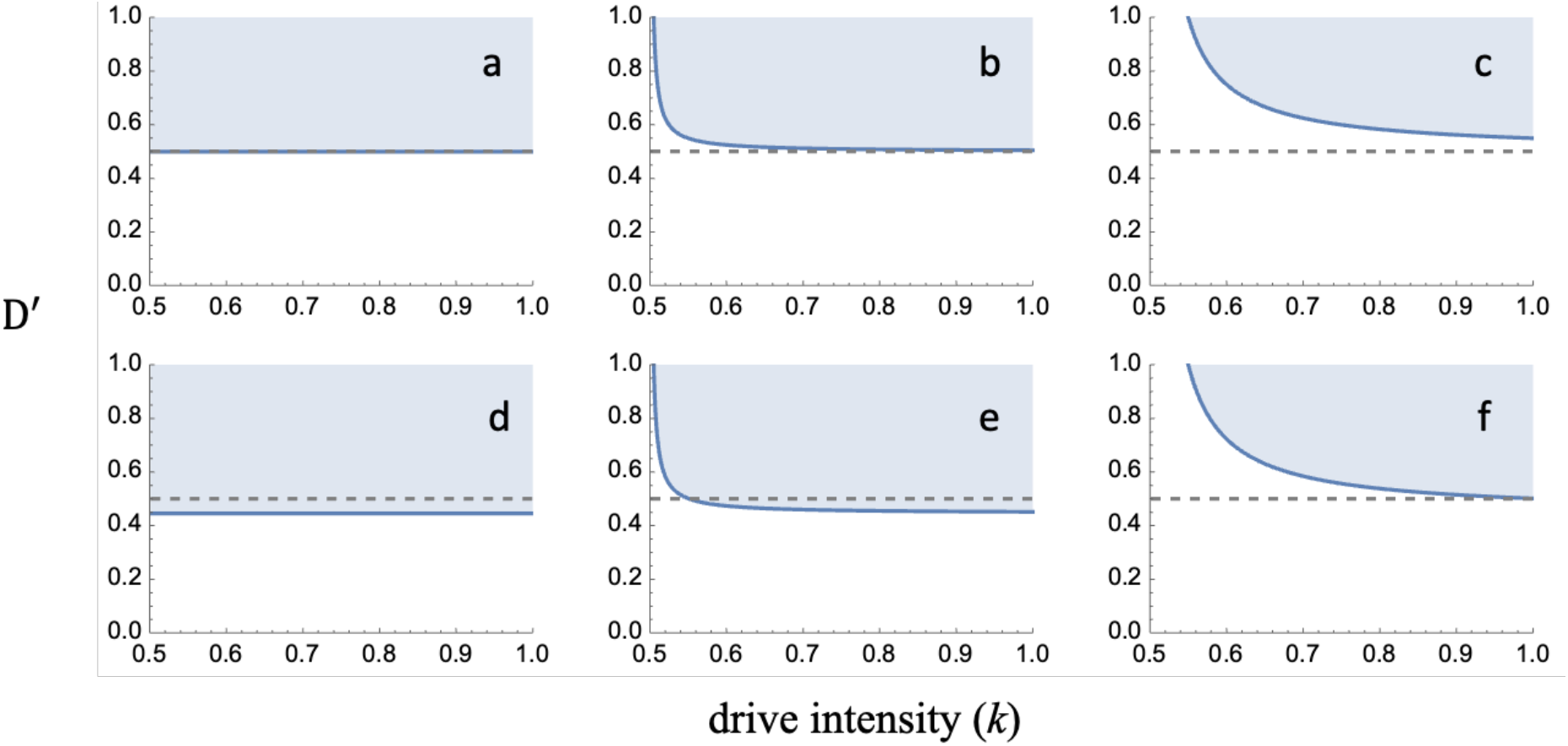
Normalized linkage disequilibrium (D’) necessary for monotonic increase of a rare centromere driver linked to a sign-changing modifier. (a-f) pA_2_ = 10^-5^; dashed line is D’ = 0.5; shaded region conveys the interval (D’_min_, 1] for the drive intensities on the *k*-axis. (a) pM_2_ = 0.0005, s = 0. (b) pM_2_ = 0.0005, s = 0.01. (c) pM_2_ = 0.0005, s = 0.1. (d) pM_2_ = 0.05, s = 0. (e) pM_2_ = 0.05, s = 0.01. (f) pM_2_ = 0.05, s = 0.1.

### 3.2 Step two: evolution of a centric fusion unlinked to the modifier

Following the fixation of A_2_, the intermediate frequency of M_2_ may make it possible for chromosomes unlinked to M to undergo karyotypic evolution by centromere drive. The equations are the same as in the **Model** section, except r = ½ is assumed. The haplotypes are now labeled as B_1_|M_1_, B_1_|M_2_, B_2_|M_1_, and B_2_|M_2_, where | indicates free recombination between loci B and M; the corresponding gametic frequencies (males, females) are x_1_, y_1_; x_2_, y_2_; x_3_, y_3_; and x_4_, y_4_, respectively (repurposing variables from **Section 3.1** and the **Model** section; *k* and *s* describe the B locus now).

The resulting Jacobian matrix is evaluated at fixation of B_1_ and a neutral posthitchhiking equilibrium of 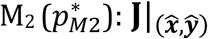, where 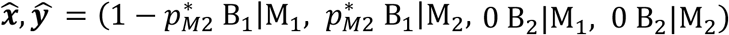. This yields a **J_mut_** submatrix with a leading eigenvalue corresponding to

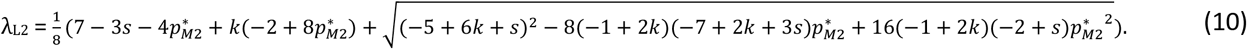

#### Result 2

|λ_L2_| > 1 when:

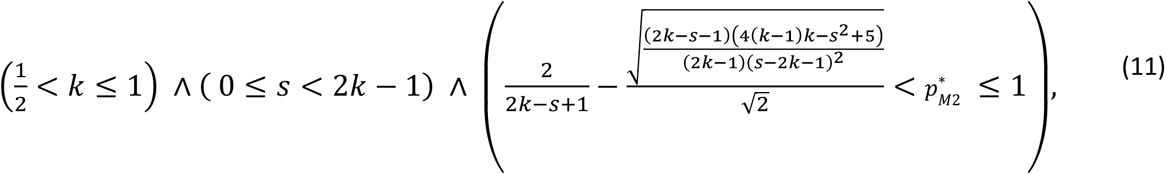

implying invasion of the unlinked B2 centric-fusion, so long as the conditions above are satisfied with respect to the parameters of drive intensity (*k*), selection (*s*), and the post-hitchhiking frequency of the dominant sign-changing modifier 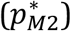.

The modifier locus does not evolve in the unlinked case. Since no recessive fitness costs are assumed at B, a system satisfying the invasion conditions of **Result 2** causes the fixation of the modifier-unlinked B_2_ centric fusion.

I define the left-hand side of the inequality involving 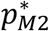 as 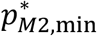 in expression (11) and this corresponds to the minimal mutant modifier frequency necessary for the spread of an unlinked centric fusion driver. A population which experiences a fixation at the modifier-linked chromosome (i.e. step 1) will undergo karyotypic changes throughout the genome if (i) the post-hitchhiking frequency of the M_2_ modifier is above 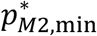 and (ii) the unlinked chromosomes have suitable selection and drive parameters that conform to the *k* and *s* inequalities above. The value of 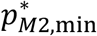 itself depends on the specific characters of a focal driver (that is, on the *k* and *s* parameters). On the simplifying assumption that karyotypic evolution is typified by fixed parameters of *s* and *k* for all drivers, then 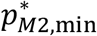 corresponds to the genome-wide minimal frequency for karyotype evolution by sign-epistatic centromere drive. A contour plot showing the parameter space associated with the step 2 invasion conditions is given in Fig. 3.

**Fig 3.**
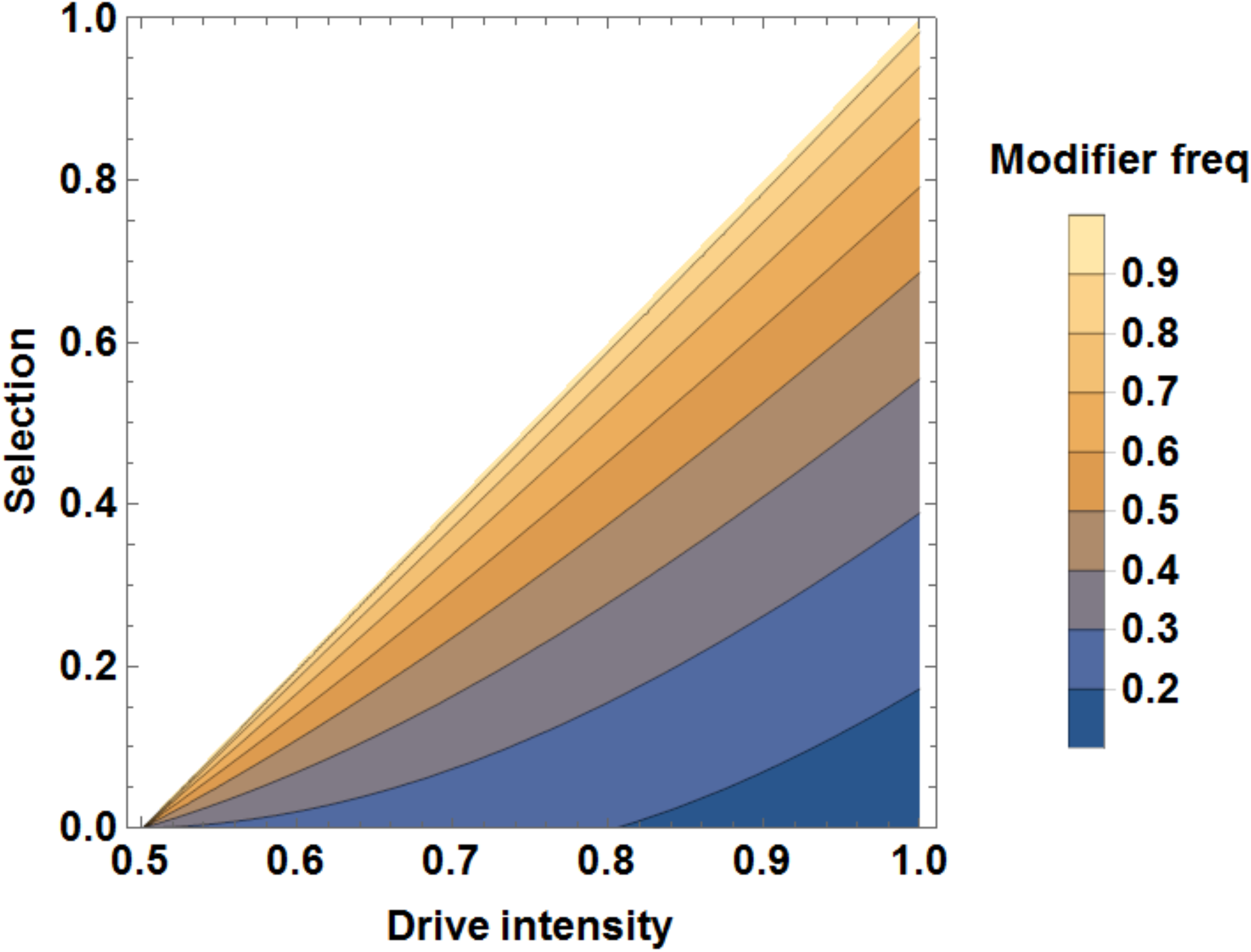
Invasible parameter region for an unlinked centric fusion, subsequent to the invasion of an unlinked modifier-centric-fusion gene complex. The space of parameters from Result 2 are displayed, with drive intensity in female meiosis on the horizontal axis, the selection coefficient of male underdominance on the vertical, and the posthitchhiking modifier frequency 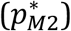 given by the contours.

A critical question remains to be answered: is 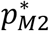 typically greater than 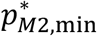? That is, does the modifier generally hitchhike to such a high frequency as to permit unlinked karyotypic evolution? Lacking an explicit derivation of the magnitude of hitchhiking, I turn to numerical iterations of the “step 1” process. The distribution of the final state of pM_2_ trajectories will be compared to that of 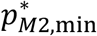 to see if the step 1 invasion is typically expected to proceed to step 2.

### 3.3 Numerical iterations of the hitchhiking effect

I obtained 10^5^ “monotonically invasible” parameter sets by sampling from the multivariate uniform distribution bounded by the parameter space in **Result 1b** (for *k*, *s*, pM_2,0_, and *r* parameters) and the corresponding linkage disequilibrium region 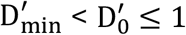. Assuming an initial pA_2_ = 10^-5^ and equality of gametic frequencies in males and females, iterations were run until Δ(pA_2_) < 10^-10^ or until 50,000 generations (whichever came first). Ignoring six cases of failed A_2_ fixation by the preceding criteria, iterations uniformly resulted in 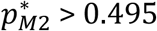. Moreover, 95% of cases yielded in 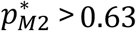; 80% of cases yielded 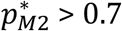; and the mean modifier outcome is 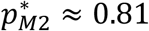 (Fig. 4a). It is apparent that, typically, only a small sliver of the depicted parameter space in Fig. 3 would restrict the evolution of an unlinked karyotypic driver. That is, assuming 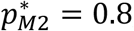, only the space spanning the two topmost contours would be impermissible to unlinked invasion. The “step 2” process is therefore well within reach of unlinked chromosome variants in the aftermath of the initial fixation event. Additionally, the time course of the step 1 fixation is rapid (mean time to A2 fixation ≈ 276 generations; Fig S2).

**Fig 4.**
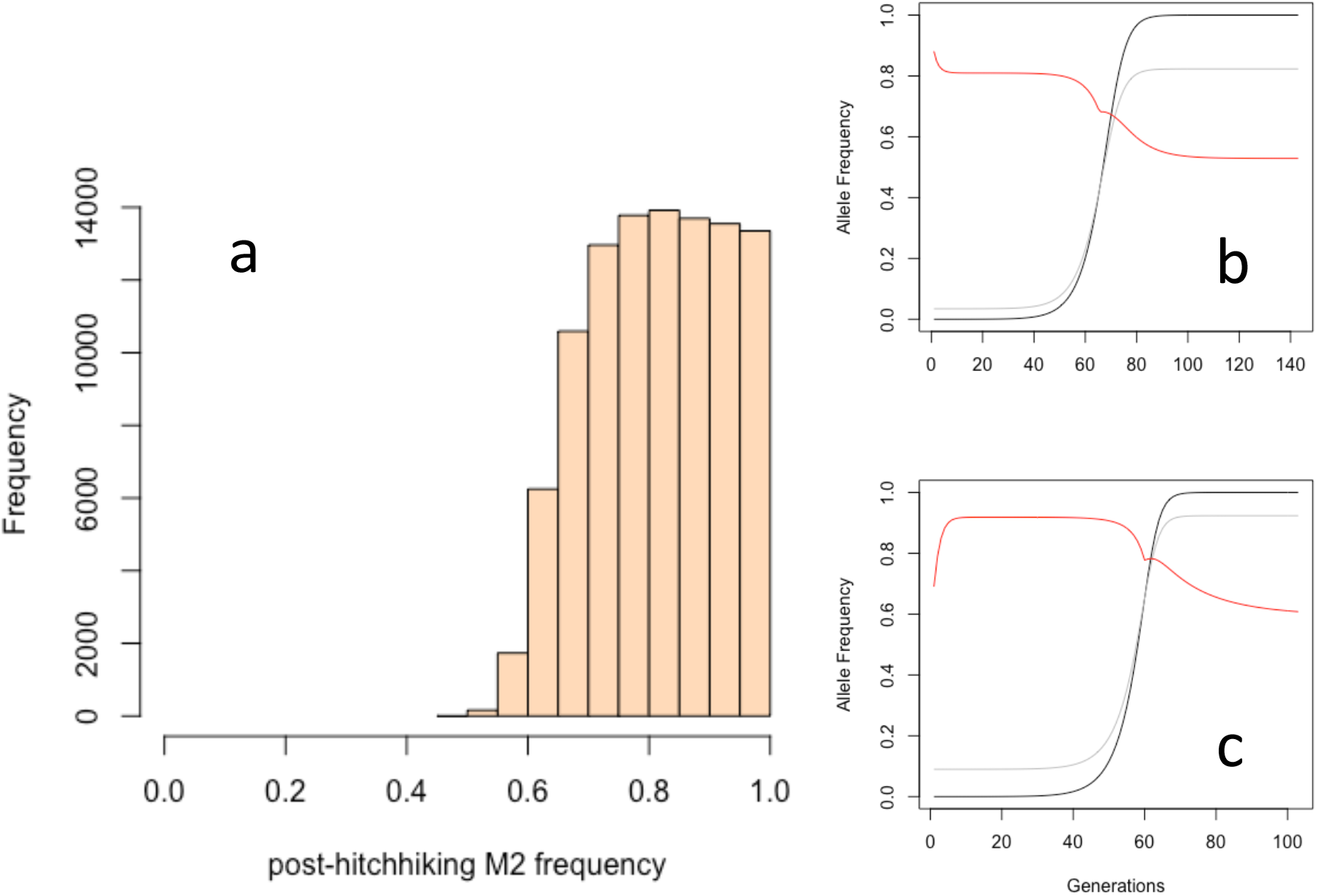
Numerical results for the modifier-linked phase of karyotype drive. (a) Histogram of post-hitchhiking M2 frequency 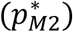 for 99,994 successful numerical runs (reached A_2_ fixation). Mean value 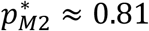. (b-c) Example trajectories: A_2_ (black), M_2_ (gray), D’ (red). (b) *k* = 0.8101498, *s* = 0.03285454, pM_2,0_ = 0.03437736, *r* = 0.08882044, 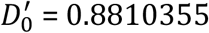. (c) *k* = 0.8566738, *s* = 0.1828839, pM_2,0_ = 0.08975939, *r* = 0.04205685, 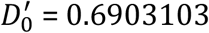.

### 3.4 Additional results

The **Appendix** derives the invasion conditions when a modifier not only changes the sign of the segregation ratio, but also alters the magnitude of drive intensity (sign-and-magnitude epistatic female drive)

## Discussion

The invasion analysis presented here provides theoretical support for Pardo-Manuel de Villena and Sapienza’s argument (2001) that the evolution of meiotic spindles brings about whole karyotype evolution.^8^ The model predicts that a modifier-linked chromosome variant will always be the first to invade, with invasion in the rest of the genome having to wait upon sufficient increase of the *trans-*acting, sign-reversing modifier allele. Numerical results indicate that the ultimate frequency of the modifier after hitchhiking along with the centric fusion is, for the vast majority of parameters, greater than the minimal frequency needed for subsequent evolutionary changes at unlinked chromosomes.

The pattern of karyotypic orthoselection^16^ for acro- and metacentrics, as revealed by the bimodality of their distribution within clades, as well as the observation of sister taxa and intraspecific demes with widely divergent karyotypes, may possibly owe to evolutionary factors other than meiotic drive, including some combination of individual-level selection, mutational biases, random drift and inbreeding.^17–19^ Strong evidence for the karyotypic drive hypothesis comes from experimental work that directly implicates the operation of strong-centromere drive in females as a causal factor in the switch from acrocentric to metacentric forms in populations of *Mus.^5^*

Although drift and inbreeding are undoubtedly relevant factors in the evolution of underdominant chromosomal rearrangements, these forces do not imply systematic homogenizing effects on the whole karyotype across taxa with diverse patterns of population structure.^18^ The operation of a sign-changing global modifier of meiotic drive offers both (1) an explanation of within-species uniformity of chromosome form as owing to meiotic drag of variants disfavored by the modifier and (2) an explanation of between-species divergence of form as owing to a co-evolutionary invasion of a karyotypic driver and a dominant modifier, followed by karyotypic invasions among the remaining chromosomes. Adding to the plausibility of the drive mechanism is that the magnitudes of distortion in known cases of centromere drive correspond to an extremely strong force of directional evolution (i.e. ~60:40 segregation). This is important as fitness costs of karyotypic heterozygosity, even in single Rb heterozygotes, can sometimes be substantial.^20^

Chmátal et al (2014) argue for the existence of population-specific correlations between chromosome form and centromere strength.^5^ Their suggestion forms a plausible alternative to the scenario investigated here in that the *correlation* is the object of reversal by genetic and evolutionary factors that affect the background centromere strength; this correlative aspect of the centric fusion/fission mutational process is proposed to be sensitive to the wild-type level of centromere strength. Such a scenario would not require the reorientation of a spindle character in order to effect a karyotypic switch, as has been modeled here. On the other hand, when Akera et al. (2017) discovered the existence of a relevant spindle asymmetry in *Mus*, they were also able to discover causal genes.^7^ Do alleles at these genes or at interacting loci ever cause the reorientation of spindle polarity? To my knowledge, this remains an open question. The differentiation of the two alternative hypotheses as a general mechanism is especially pertinent to the following: is it ever the case that *weaker* centromeres benefit from preferential capture by the egg pole? The flipping of spindle-polarity entails that whatever drives on the old spindle will drag on the reversed spindle, whether this be on the basis of differential centromere number or differential centromere strength. A stronger-always-wins hypothesis necessarily accepts the existence of an asymmetric spindle but militates against its reorientation over evolutionary time, as a reversal would cause weaker centromeres to be favored in some lineages.

It remains to ask why spindles are asymmetric with respect to their microtubule organization in the first place. The adaptive benefit of such differences across the spindle is not clear (but see Ubeda and Haig 2005, Brud 2022). In the absence of an adaptive purpose for the trait, presumably female drive suppressors have periodically abolished microtubule differences across the spindle, only to have them restored by asymmetry-enhancers that benefit from hitchhiking with centromeres. The evolutionary struggle between such genes seems to have been won decisively by the asymmetry-enhancers, as we can deduce from the observation of female drive in diverse taxa.^3^ Although the reason for the victory is unclear, Crow (1991) argued that drive-enhancers tend to spread more quickly than drive-suppressors, and so “a few linked enhancers may sometimes outweigh a larger number of suppressors”.^21^ If that’s the case, then fundamental features of meiosis may reflect the anti-adaptationism of selfish genetic elements.^22,23^

## Supporting information

Mathematica nb, R script, and parameter file

---

**Fig. S1.**
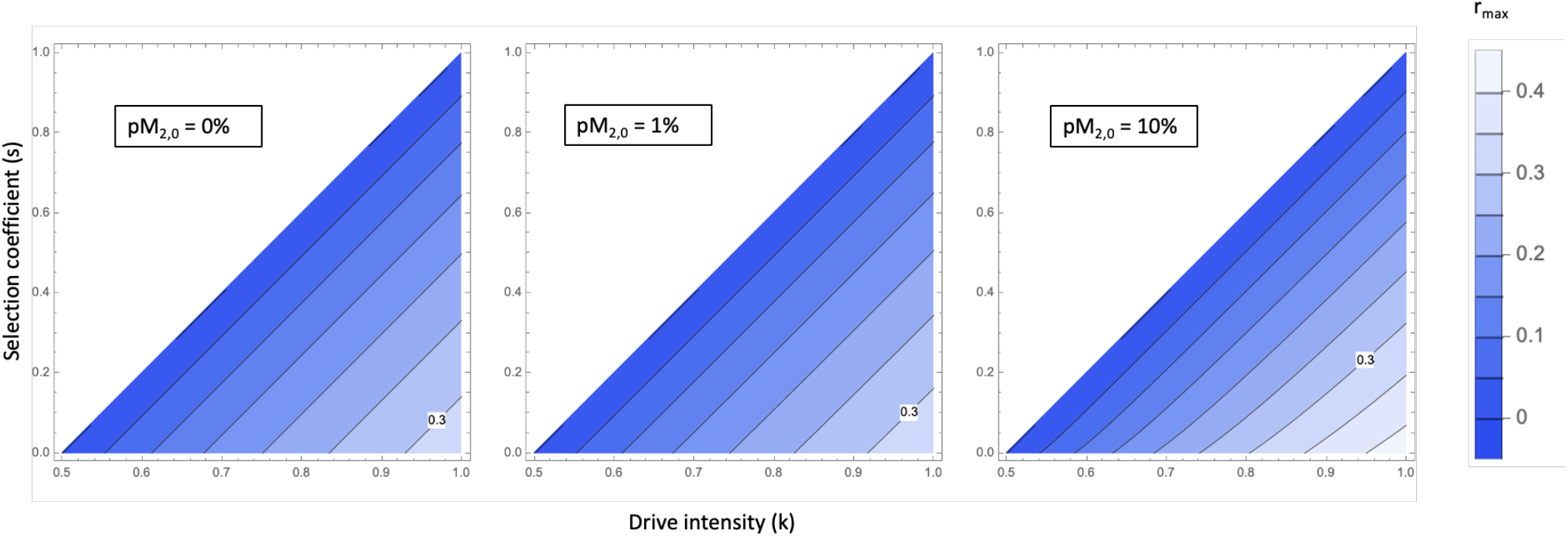
Parameter-spaces permitting double-invasion. Contours indicate the value r_max_ consistent with the drive intensity and selection coefficient given on the k and s axes, respectively, and demarcate changes of 0.05. The initial frequency of M_2_ is framed in each panel. The value r_max_ = 0.3 is labeled for comparison.

**Fig. S2.**
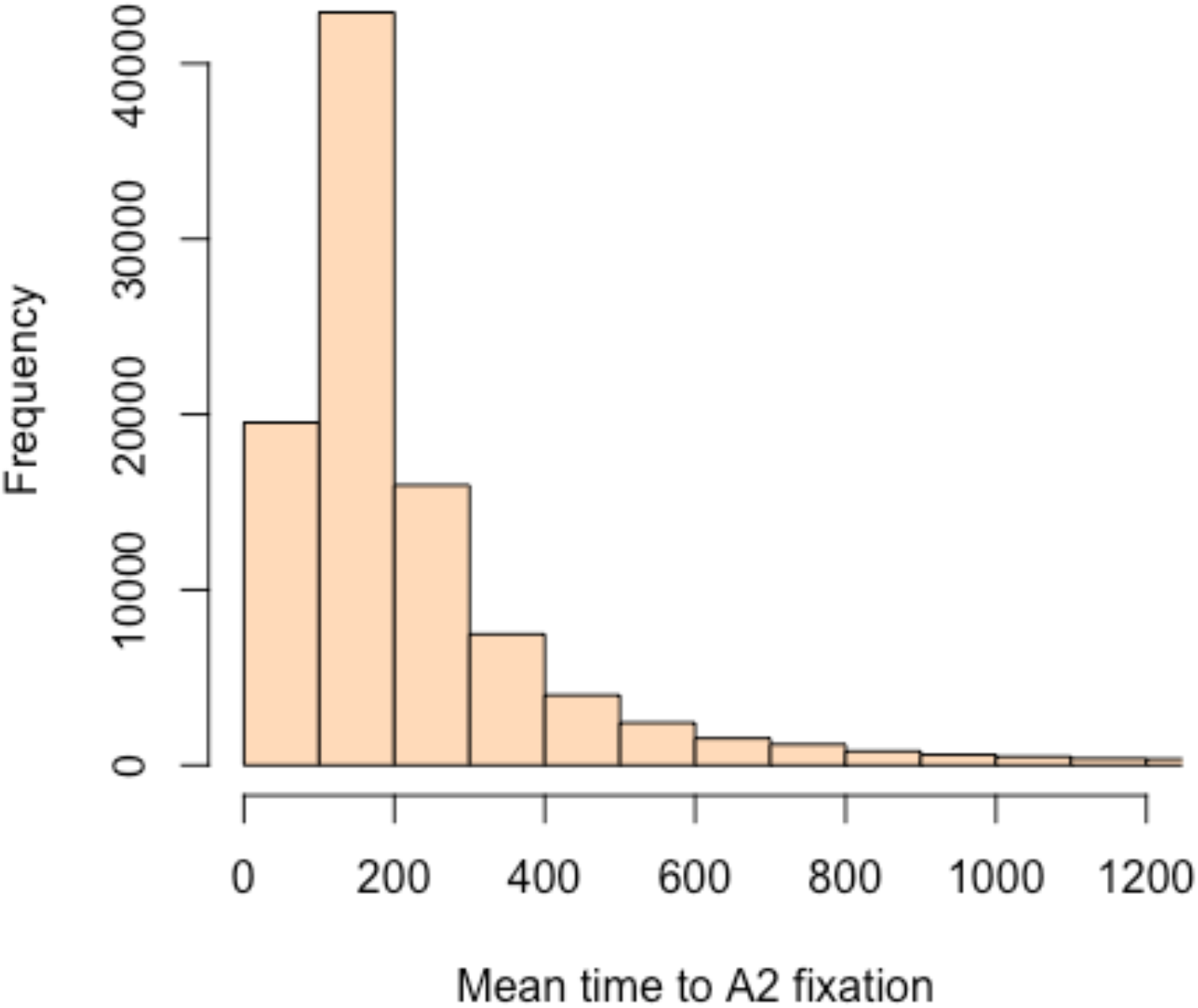
Mean time to fixation of A_2_ (modifier-linked centric fusion driver). Histogram of 99,994 successful A_2_ fixations (as declared when Δ(pA_2_) < 10^-10^; A_2_ trajectories uniformly attained “1.0” numerical values in R for these cases). Mean time to fixation is approx. 276 generations. Rarer bins on the x-axis (to the right of 1200 generations) not shown.

## Appendix. Sign-and-magnitude epistatic centromere drive

Assume the following female segregation matrix **K**, featuring a resident drive-intensity *k*_1_ and a mutant drive intensity *k*_2_. That is, *k*_1_ and *k*_2_ are different magnitudes, but both lie in the interval [0.5, 1]. The M_2_ modifier flips the direction of the segregation advantage and simultaneously enacts the *k*_2_ intensity favoring A_2_. Here it is:

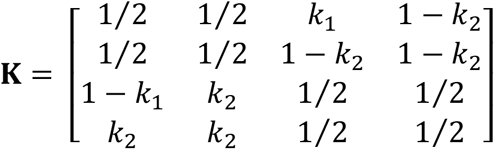

The **Model** section is otherwise unchanged except: *c*_1_ = – *k*_2_), *c*_1_ = 1 – *k*_2_, *c*_3_ = *k*_2_, and *c*_4_ = (–1)*k*_2_.

Step 1:

The leading eigenvalue associated with double-invasion (from 100% A_1_M_1_ resident population and 0% for the remaining haplotypes):

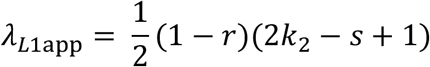

Modifier linked invasion conditions:

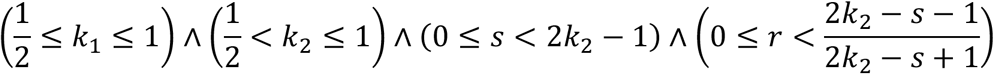

Step 2:

The leading eigenvalue associated with subsequent evolution of unlinked karyotypic driver (from 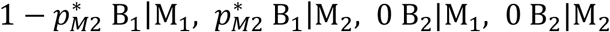):

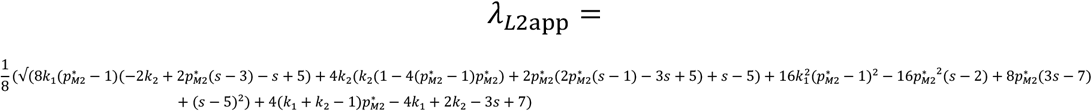

Unlinked invasion conditions:

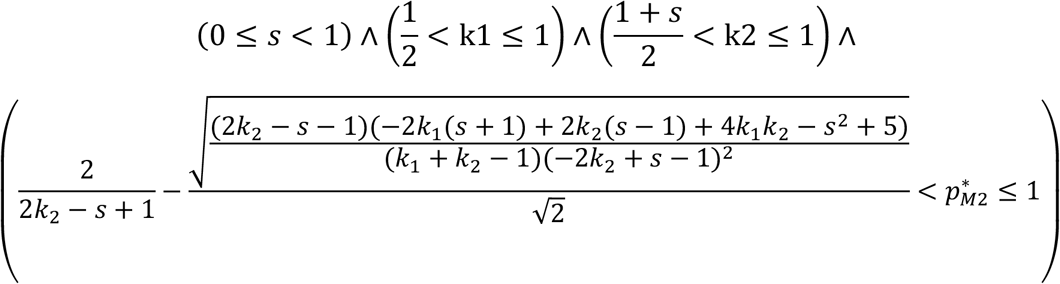

