## Supplementary material for "Sign-epistatic centromere drive in panmictic populations": Mathematica nb, R script, and parameter file: SignEpi_Drive.R

```
SexLim.Sign.Epi.Drive <- function(k,s,pm0,r,Dprime){
  
  f.A2 <- 0.00001
  
  f.M2 <- pm0
  
  f.A1 <- 1-f.A2
  f.M1 <- 1-f.M2
  
   Dmax <- min(f.A1*f.M2, f.A2*f.M1)
  # D <- min(f.A1*f.M2, f.A2*f.M1)
   
   D <- Dprime*Dmax
  
  x1 <- f.A1*f.M1 + D   # A1M1
  x2 <- f.A1*f.M2 - D  # A1M2
  x3 <- f.A2*f.M1 - D  # A2M1
  x4 <- f.A2*f.M2 + D  # A2M2
  
  y1 <- f.A1*f.M1 + D
  y2 <- f.A1*f.M2 - D
  y3 <- f.A2*f.M1 - D
  y4 <- f.A2*f.M2 + D
  
  p1 <- (1/2)*sum(x1,y1)
  p2 <- (1/2)*sum(x2,y2)
  p3 <- (1/2)*sum(x3,y3)
  p4 <- (1/2)*sum(x4,y4)
  


  
  A2.history <- c(f.A2)
  M2.history <- c(f.M2)
  #W.history <- c()
  Dprime.history <- c(Dprime)
  D.history <- c(D)
  gen <- 0
  
  ### The value of deltaA2 below is arbitrary, used to initialize the while function that follows
  deltaA2 = 0.00001
  
  while(abs(deltaA2) > 1e-10 && gen < 50000){
    
    W <- x1*y1*(1) + x1*y2*(1) + x1*y3*(1 - s) + x1*y4*(1 - s) + x2*y1*(1) + x2*y2*(1) + x2*y3*(1 - s) + x2*y4*(1 - s) +x3*y1*(1 - s) + x3*y2*(1 - s) + x3*y3*(1) + x3*y4*(1) +x4*y1*(1 - s) + x4*y2*(1 - s) + x4*y3*(1) + x4*y4*(1)
    
    #W.history <- append(W.history,W)
    
    ### Evolution of male haplotype frequencies
    
    x1.prime <- ((1/W)*(x1*(y1*(1/2) + y2*(1/2) + y3*(1/2)*(1 - s) + y4*(1/2)*(1 - s)) + y1*(x1*(1/2) + x2*(1/2) + x3*(1/2)*(1 - s) + x4*(1/2)*(1 - s)) - (r)*(1/2)*(1 - s)*(x1*y4 + x4*y1 - x2*y3 - x3*y2)))
    
    x2.prime <- ((1/W)*(x2*(y1*(1/2) + y2*(1/2) + y3*(1/2)*(1 - s) + y4*(1/2)*(1 - s)) + y2*(x1*(1/2) + x2*(1/2) + x3*(1/2)*(1 - s) + x4*(1/2)*(1 - s)) + (r)*(1/2)*(1 - s)*(x1*y4 + x4*y1 - x2*y3 - x3*y2)))
    
    x3.prime <- ((1/W)*(x3*(y1*(1/2)*(1 - s) + y2*(1/2)*(1 - s) + y3*(1/2) + y4*(1/2)) + y3*(x1*(1/2)*(1 - s) + x2*(1/2)*(1 - s) + x3*(1/2) + x4*(1/2)) + (r)*(1/2)*(1 - s)*(x1*y4 + x4*y1 - x2*y3 - x3*y2)))

    x4.prime <- ((1/W)*(x4*(y1*(1/2)*(1 - s) + y2*(1/2)*(1 - s) + y3*(1/2) + y4*(1/2))+ y4*(x1*(1/2)*(1 - s) + x2*(1/2)*(1 - s) + x3*(1/2) + x4*(1/2)) - (r)*(1/2)*(1 - s)*(x1*y4 + x4*y1 - x2*y3 - x3*y2)))
    
    ### Evolution of female haplotype frequencies
    
    y1.prime <-        x1*(y1*(1/2) + y2*(1/2) + y3*(k) + y4*(1 - k)) + y1*(x1*(1/2) + x2*(1/2) + x3*(k) + x4*(1 - k)) - (r)*(1 - k)*(x1*y4 + x4*y1 - x2*y3 - x3*y2)

    y2.prime <-        x2*(y1*(1/2) + y2*(1/2) + y3*(1 - k) + y4*(1 - k))+ y2*(x1*(1/2) + x2*(1/2) + x3*(1 - k) + x4*(1 - k)) + (r)*(1 - k)*(x1*y4 + x4*y1 - x2*y3 - x3*y2)
  
    y3.prime <-        x3*(y1*(1 - k) +  y2*(k) + y3*(1/2) + y4*(1/2)) + y3*(x1*(1 - k) +  x2*(k) + x3*(1/2) + x4*(1/2)) + (r)*(k)*(x1*y4 + x4*y1 - x2*y3 - x3*y2)
    
    y4.prime <-        x4*(y1*(k) +  y2*(k) + y3*(1/2) + y4*(1/2)) + y4*(x1*(k) +  x2*(k) + x3*(1/2) + x4*(1/2)) - (r)*(k)*(x1*y4 + x4*y1 - x2*y3 - x3*y2)
    

    #####
    
    
    x1 <- x1.prime
    x2 <- x2.prime
    x3 <- x3.prime
    x4 <- x4.prime
    
    y1 <- y1.prime
    y2 <- y2.prime
    y3 <- y3.prime
    y4 <- y4.prime
    
    
    p1 <- (1/2)*sum(x1,y1)
    p2 <- (1/2)*sum(x2,y2)
    p3 <- (1/2)*sum(x3,y3)
    p4 <- (1/2)*sum(x4,y4)
    
    f.A1 <- p1+p2
    f.A2 <- p3+p4
    
    f.M1 <- p1+p3
    f.M2 <- p2+p4
    
    
    D <- p4 - f.A2*f.M2
    
    
    if(D >= 0){Dprime <- D/min(f.A1*f.M2, f.A2*f.M1)}   ### D.prime (Lewontin's D) calculated from sex-averaged allele frequencies
    if(D < 0 ){Dprime <- D/min(f.A1*f.M1, f.A2*f.M2)}

    
    A2.history <- append(A2.history, f.A2)
    
  
    M2.history <- append(M2.history, f.M2)
    
    
    Dprime.history <- append(Dprime.history, Dprime)
    D.history <- append(D.history, D)
    
    gen <- gen+1
    deltaA2 = f.A2 - A2.history[gen]
    
   
  }
  
  plot(A2.history,type ="l",ylim = c(0,1), lwd =1, xlab = "Generations", ylab = "Allele Frequency")
  lines(M2.history, col ="grey", lwd = 1)
 lines(Dprime.history, col = "red")
#  lines(D.history*4, col = "blue")
#  print(head(A2.history,50))
  
#print(c(tail(M2.history,1),tail(A2.history,1), gen))
#   plot(M2.history)

 
}
```
